# 3DPolyS-LE: an accessible simulation framework to model the interplay between chromatin and loop extrusion

**DOI:** 10.1101/2022.04.15.488456

**Authors:** Todor Gitchev, Gabriel Zala, Peter Meister, Daniel Jost

## Abstract

**Motivation:** Recent studies suggest that the loop extrusion activity of Structural Maintenance of Chromosomes complexes is central to proper organization of genomes *in vivo*. Polymer physics-based modeling of chromosome structure has been instrumental to assess which structures such extrusion can create. Only few laboratories however have the technical and computational expertise to create *in silico* models combining dynamic features of chromatin and loop extruders.

**Results:** Here we present 3DPolyS-LE, a self-contained, easy to use modeling and simulation framework allowing non-specialists to ask how specific properties of loop extruders and boundary elements impact on 3D chromosome structure. 3DPolyS-LE also provides algorithms to compare predictions with experimental Hi-C data.

**Availability and implementation:** Software available at https://gitlab.com/togop/3DPolyS-LE ; implemented in Python and Fortran 2003 and supported on any Unix-based operating system (Linux, Mac OS).

**Supplementary Information:** Supplemental data are available at Bioinformatics online

## 1. Introduction

Genes are regulated at many levels, from local transcription factor binding to the megabase-range contacts between enhancers and promoters. Recent findings have highlighted the function of the chromosome 3D organization in the latter regulation, as the genome is partitioned into consecutive regions of enhanced compaction, the so-called ‘topologically associated domains’ (TADs), where promoters and enhancers colocalize (Dixon *et al*., 2012; Nora *et al*., 2012). At this scale, genome folding is mostly a consequence of the interplay between loop extrusion factors of the Structural Maintenance of Chromosome (SMC) complexes family and oriented boundary elements bound by proteins that limit loop extrusion (reviewed in Mirny and Dekker, 2021). In particular, by comparing results from *in silico* models and *in vivo* Hi-C data, polymer simulations proved very useful to understand the TAD structure of chromosomes, suggest and test hypotheses on the function of boundary elements or loop extrusion factors (Rao *et al*., 2017; Fudenberg *et al*., 2016; Schwarzer *et al*., 2017; Goloborodko *et al*., 2016; Brandão *et al*., 2021; Nuebler *et al*., 2018). As the development of such simulations is technically difficult and thus not accessible to biologists aiming to (in)validate a mechanistic hypothesis on TAD formation for their system of interest, we provide an open-access, user-friendly, generic modeling framework for physics-based polymer simulations of loop extrusion (3DPolyS-LE), wrapped as a Python package, allowing users to run simulations by varying parameters on boundary elements and loop extruders properties and assess the expected structures.

## 2. Model

3DPolyS-LE simulates the dynamics of one chromosome, modeled as a coarse-grain polymeric chain in which each monomer, of size 50 nm, contains 2 kb of chromatinized DNA. In absence of loop extrusion, the dynamics of the chain is governed by the generic properties of a homopolymer: chain connectivity, excluded volume and bending rigidity *(Ghosh and Jost, 2018)*. Additionally, the polymer can be extruded by loop extruding factors (LEFs) that dynamically bind and unbind from chromatin (**Fig.1A**, for details see **Supplementary Methods**). Initial binding of LEFs could be at predefined loading sites or non-specifically along the chromosome. Bound LEFs are composed of two ‘legs’ that may translocate along the genome, creating dynamic loops between gradually more distant regions along the chain. We implemented two scenarios for the leg motion: (1) symmetric extrusion with LEF legs progressing along chromatin in opposite directions at the same speed, as observed *in vitro* for cohesin (Davidson *et al*., 2019; Kim *et al*., 2019); (2) asymmetric extrusion with only one translocating leg, as observed *in vitro* for condensin (Ganji *et al*., 2018; Kong *et al*., 2020). The motion of a LEF can be restricted by the presence of boundary elements that may stop or slow down the progression of legs depending on their directionality, and by collisions with the other extruding LEFs. We integrated two scenarios for collisions between extruding legs: (1) legs are impenetrable obstacles and they cannot move until one detaches from chromatin as usually assumed for cohesin-mediated extrusion (Fudenberg *et al*., 2016); (2) legs are phantom obstacles and can cross each other. This is the so-called Z-loop process recently observed *in vitro* for yeast condensin (Kim *et al*., 2020) and *in vivo* for bacterial SMCs (Brandão *et al*., 2021).

**Figure 1.**
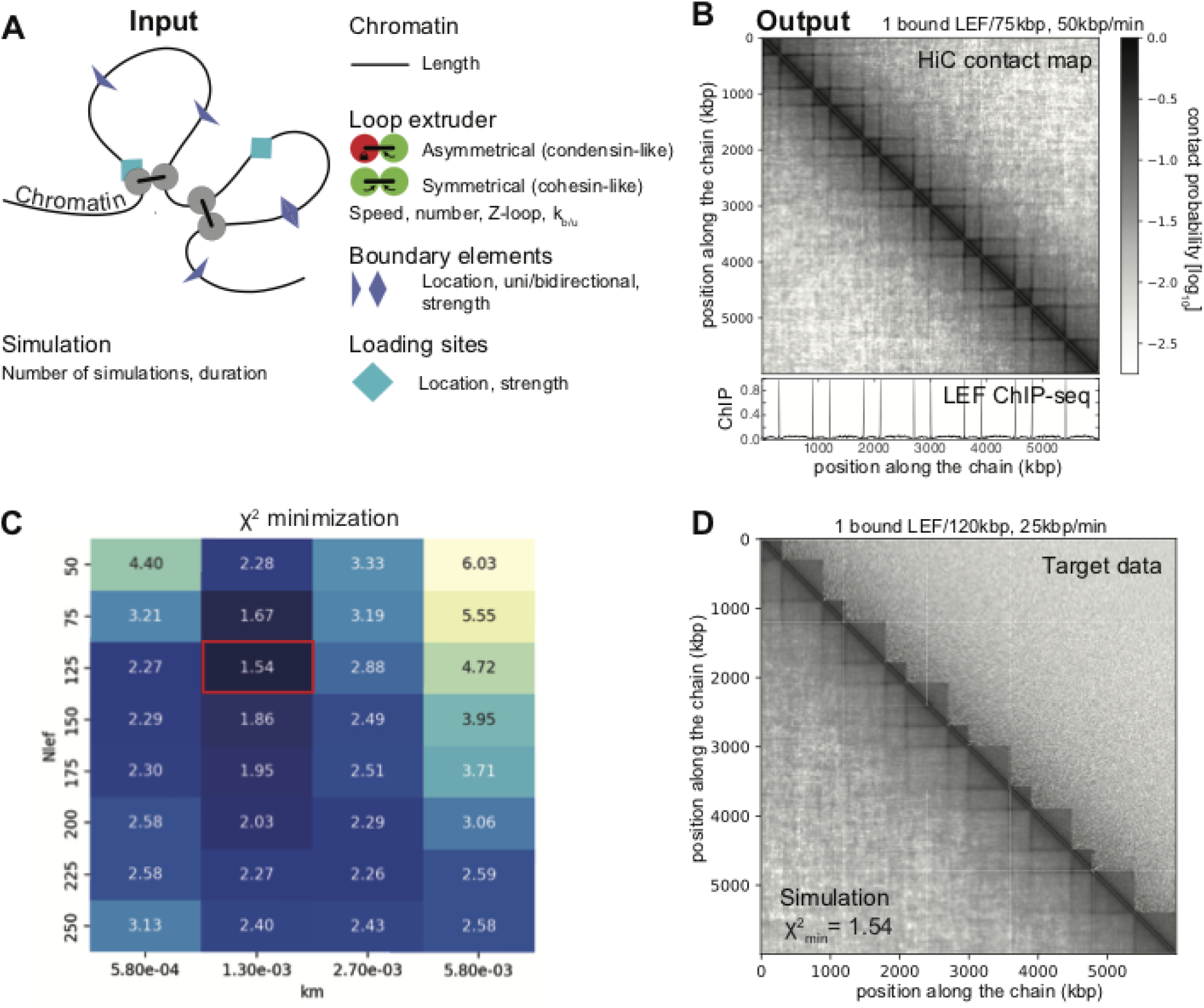
Features provided by 3DPolyS-LE. (A) Polymer model and input parameters and data. (B) Typical outputs of the simulations: virtual Hi-C data (top) and ChIP-Seq profile (bottom) of loop extruders. (C) Grid-simulations to find the best set of parameters that fit the target data. For each parameter set, a χ^2^-score is estimated. Example of optimization with synthetic human Hi-C data as target (see **Supplementary Methods**). (D) Best model predictions from the simulations in (C) (lower part) compared to target data (upper part).

## 3. Description of the program

### 3.1 Inputs and outputs

3DPolyS-LE allows to modify several parameters controlling LEF properties (binding, density, velocity), to select the leg motion type (symmetric/asymmetric) and head-to-head collisions scenarios (impenetrable/phantom), as well as to position of boundary elements (with individual directionality and strength; for a detailed description of parameters and how to change them, see **Supplementary Methods**). Depending on the model parameters and extrusion properties, 3DPolyS-LE simulates, for a given number of independent polymers, the dynamics of the chromosome during a user-defined time period. During the simulations, snapshots of the current polymer conformations and LEF positions are stored at regularly-spaced time intervals. From these snapshots, virtual Hi-C maps and ChIP-seq profiles for LEF occupancy are produced in HDF5 (with an included converted to ‘cooler’ format, (Abdennur and Mirny, 2019)) and bedGraph formats, respectively (**Fig.1B**). Optionally, three-way chromatin contacts (see **Supplementary Methods** and **Sup. Fig. 2**) can be extracted for downstream comparative analysis with data from GAM (Beagrie *et al*., 2017) or multi-contact nanopore-derived ‘C’ technologies (Allahyar *et al*., 2018; Ulahannan *et al*., 2019). If a reference Hi-C map is provided (ie.g. experimental *in vivo* data), 3DPolyS-LE will compute relevant metrics (see **Supplementary Methods**) to quantitatively compare model predictions with this data.

### 3.2 Implementation and performance

The program is organized as a Python package, requiring specific libraries for compilation and parallel processing of the core simulation module and for the downstream analysis (see **Supplementary Methods** for more details). The first step ‘Simulations’ is running simulations defined by the configuration files describing parameter values, with the possibility to use multiple cores of a High Performance Computing (HPC) cluster over a Message Passing Interface (MPI) framework. The next step called ‘Analysis’ processes simulation results and extracts Hi-C and ChIP-seq data. The last step ‘Comparison’ is the comparison to a provided reference dataset and the production of related plots. In the case of a series of simulations with different parameters (or ‘grid’-simulations, see **Supplementary Methods**), summary statistics can be visualized in a heat-map plot. All steps are run with a single command using a scheduler. The package is working on any Unix-based operating system (Linux, MacOS) and has been tested on a Slurm HPC cluster. Using a polymer equivalent to a 6 Mb chromosome (3000 beads), a two hours (real time) simulation required roughly 20 CPU.min (AMD Epyc, 4 Gb RAM).

### 3.3 Examples

As an illustration of 3DPolyS-LE, we modeled loop extrusion by cohesins during interphase in mammals, by simulating a 6 Mb-long polymer with impermeable boundaries placed along the chain every 300 kb or 600 kb with symmetric leg motion, impenetrable head-to-head collisions, random loading onto the polymer and default binding/unbinding rates estimated from *in vivo* imaging data (see **Supplementary Methods**) (Cattoglio *et al*., 2019; Hansen *et al*., 2017). For a density of 1 bound LEF per 75 kb and an extruding velocity of 50 kb/min, we observed the formation of TADs with corner peaks (**Fig.1B**). We then varied systematically the density of bound LEFs and their extruding velocity and compared the predicted intra-TAD contact probabilities to the corresponding quantity extracted from GM12878 Hi-C data for TADs of the same size (**Fig.1C,D**) (Rao *et al*., 2014). We found an optimal set of parameters of 1 bound LEF per 120 kb and a velocity of 25 kb/min, in the good range of *in vivo* (1 bound LEF every 186-372 kb(Cattoglio *et al*., 2019)) and *in vitro* estimations (30-120 kb/min,(Golfier *et al*., 2020; Kim *et al*., 2019; Davidson *et al*., 2019)), respectively. Other examples showing the impact of different scenarios (asymmetric leg motion, phantom collisions, loading at specific sites, boundary directionality, permeable boundaries, etc.) are given in **Supplementary Figure 1**.

## Funding

DJ acknowledges Agence Nationale de la Recherche [ANR-18-CE12-0006-03, ANR-18-CE45-0022-01] for funding. The PM laboratory is funded by SNF 31003A_176226 and the University of Bern. This work was made possible by the COST Action CA18127 “INC”.

## Notes

https://gitlab.com/togop/3DPolyS-LE

### Competing Interest Statement

The authors have declared no competing interest.

