## Supplementary Methods for "3DPolyS-LE: an accessible simulation framework to model the interplay between chromatin and loop extrusion"

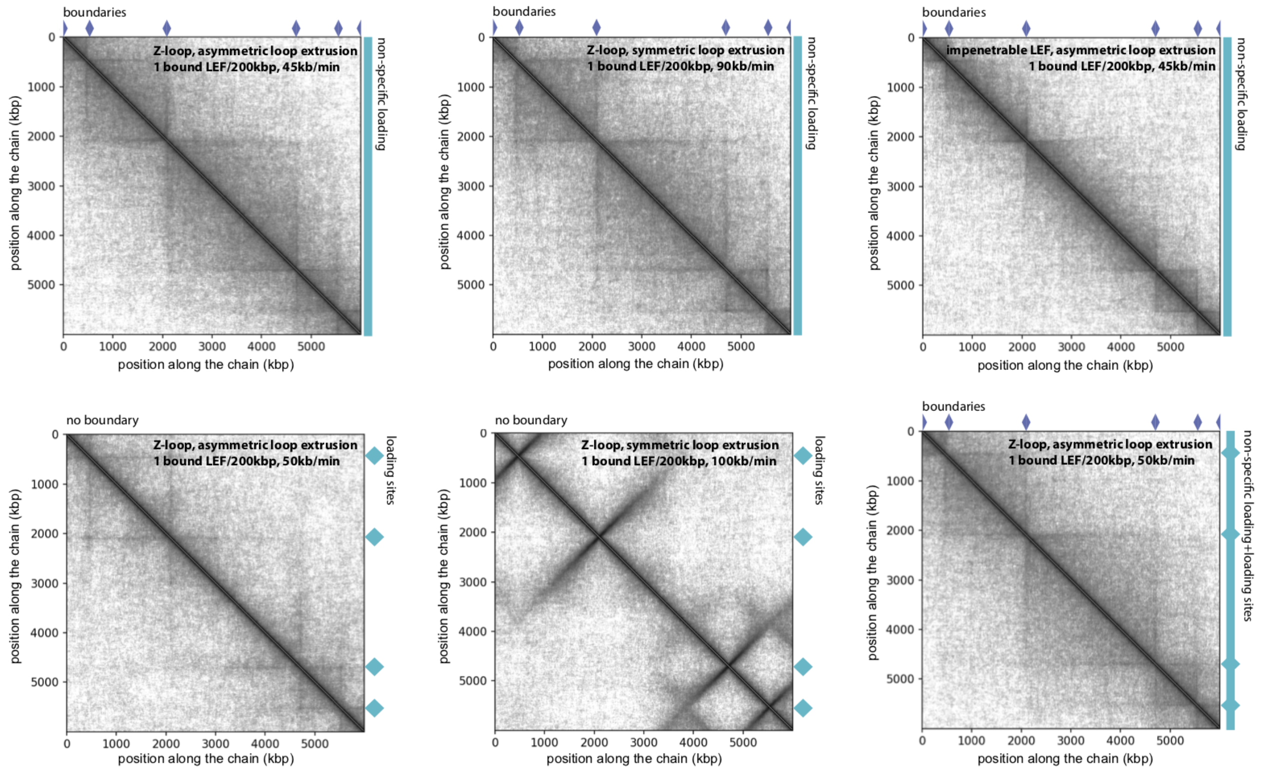


**Supplementary Figure 1.** Examples of simulated Hi-C maps for several loop extrusion scenarios.

### Polymer model

One chromosome is modeled as a semi-flexible, self-avoiding polymeric chain of $N_{chain}$ monomers; each, of diameter 50 nm, containing 2 kb of genome. The chain configurations are sampled on a FCC lattice using periodic boundary conditions to effectively account for nuclear crowding by other chromosomes, as described in (Ghosh and Jost, 2018) .

In absence of loop extrusion, the Hamiltonian of a configuration is given by $H = \kappa \sum_{i=2}^{N-1} (1 - \cos cos\, \theta_i)$, with $\kappa$ the bending rigidity and $\theta_i$ the bending angle between monomers *i-1, i* and *i+1*. The lattice volumic fraction (~0.5) and bending energy ($\kappa$ =1.17 kT) were fixed to simulate the coarse-grained dynamics of a chromatin fiber of Kuhn length ~100 nm (Socol *et al.*, 2019; Arbona *et al.*, 2017) and of a typical bp-density ~0.01 bp/nm3. Dynamics of the chain was simulated using kinetic Monte-Carlo, starting from unknotted, compact configurations, as described in (Ghosh and Jost, 2018). Each Monte-Carlo time step (MCS) consists of $N_{chain}$ local trial moves. Time mapping between simulations (MCS) and real time is performed by mapping predicted and observed mean-squared displacement as described in (Ghosh and Jost, 2018; Salari *et al.*, 2022) . At our resolution, we found 1MCS ≃ 5 msec.

To model loop extrusion, we considered, in addition, $N_{LEF}$ loop extruders that may bind to chromatin at position *i* with rate $k_b(i)$ or unbind at a position-independent rate $2k_u$. Each bound extruder has two legs that are localized on different monomers (initially nearest-neighbor along the chain at the moment of the binding). In the symmetrical scenario, the two legs 'walk' on opposite directions along the polymer, each at a rate $k_m(i, s)$ with $i$ the position along the chain of the leg and $s = \pm 1$ the direction of the leg motion (+1 if this is a leg going from i to i+1; -1 if it goes from i to i-1). In the asymmetrical scenario, only one, randomly-chosen, leg may move with rate $k_m(i, s)$, the other remaining fixed during the whole lifetime of the LEF onto chromatin. Impermeable boundaries in one direction $s$ would thus correspond in both scenarios to $k_m(i, s) = 0$. Regarding the unidimensional traffic of LEF legs along the chain, we allowed two options: (1) one monomer can be occupied by only one leg, and thus legs are impenetrable obstacles and cannot cross each other; (2) there is no limitation in monomer occupancy and thus legs are phantom obstacles and can cross each other (Z-loop scenario). Contrary to previous implementations of the loop extrusion model (Salari *et al.*, 2022; Fudenberg *et al.*, 2016; Sanborn *et al.*, 2015; Ghosh and Jost, 2020) where the LEF 1D motion were independent of the local 3D chromatin organization, we assumed here that one leg (on monomer *i*, at 3D position $\vec{x}_i$) can move (to monomer *i*+*s*) if and only if it remains in the spatial vicinity of the other leg (on monomer *j* at 3D position $\vec{x}_j$) forming the same LEF, ie if $\vec{x}_{i+s}$ is still nearest-neighbor (on the 3D lattice) of $\vec{x}_j$. This may more realistically account for the stalling of LEF motion observed *in vitro* for locally stretched chromatin configuration (Golfier *et al.*, 2020). Similarly, monomer trial moves (see above) that do not conserve the spatial proximity of legs of the same LEF are rejected. In our kinetic Monte-Carlo framework, at every MCS, $N_{LEF}$ trial attempts to bind or unbind extruders and $2N_{LEF}$ trial attempts to move a leg of a bound extruder are also performed. Overall the complexity of the algorithm is O($N_{chain}$+$N_{LEF}$).

Default parameter value for $k_u$ was fixed to fit the experimentally-observed life time of bound cohesin on chromatin (~20 min) (Hansen *et al.*, 2017) and for $k_b(i)$ such that the average proportion of bound extruders $< k_b > /(< k_b >+ 2k_u)$ is about 40% (Cattoglio *et al.*, 2019) with $< k_b > = \left[\frac{1}{N_{chain}} \sum_i k_b(i)\right]$ the average binding rate along the chain.

### Numerical simulations and parameters

The polymer model has been wrapped into a user-friendly Python package (3DpolyS-LE) where all parameters can be easily modified (see below for a technical description of the package and descriptions of input and output files). The core simulation module is written in Fortran and can be parallelized in HPC clusters. The Python interface runs the simulations and performs initial statistical analysis of the results. It is freely available on https://gitlab.com/togop/3DPolyS-LE.

#### Input parameters

A typical simulation scenario would have to define the polymer parameters (size of the chain, density, rigidity), loop extrusion parameters (velocity, occupancy of LEFs, leg motion type, head-to-head collisions scenarios, position and strength of boundary elements), simulation parameters (duration of one trajectory, number of trajectory, number of measurements along the trajectory) and analysis parameters (radius of contact).

For polymer parameters:

- *Nchain [integer*$\geq 1$*]* defines the number of monomers of the chain. *Nchain x 2kb* represents the length of the polymer in bp.
- *L [integer*$\geq 1$*]* defines the length of the box in which the polymer is simulated. It should be chosen such that $N_{chain}/(4L^3) \simeq 0.5$ to fix the bp volumic density to ~0.01 bp/nm3.
- *kint [real*$\geq 0$*]* defines the binding energy in kT-unit. It should remain fixed to 1.17.

For loop extrusion parameters:

- *km [1≥real*$\geq 0$*]* defines the maximal motion rate of one LEF leg and is given in monomer/MCS unit. To translate into kb/min: 1 kb/min corresponds to *km=1.08e-4*. Note that the actual loop extrusion speed is given by $n_{leg} \times km$ with $n_{leg}$ = 1 or 2 is the number of translocating legs (1 for asymmetric, 2 for symmetric).

- *ku [1≥real≥ 0]* defines half of the LEF unbinding rate. *ku=2e-6* (default value) corresponds to an average dwell-time of LEF on chromatin of 20 min. *ku=4e-6* to 10 min.
- *kb [1≥real≥ 0]* defines the maximal LEF binding rate. It fixes the proportion of LEF that are bound to chromatin. *kb=2.8e-6* (default value) correspond to $kb/(kb + 2ku) \simeq 40\%$ of bound LEFs (for *ku* default value). *kb=6e-6* to 60%.
- *Nlef [integer≥ 0]* defines the total number of LEFs. The density of bound LEFs (in unit of number of LEF per kb) is given by $kb/(kb + 2ku) \times N_{lef}/(N_{chain} \times 2)$. For example, for *Nchain=10,000* and default parameter for *ku,kb*, *Nlef=500* corresponds to 1 bound LEF every 100 kb.
- *lef_loading_site* specifies the file path to a CSV (comma-separated values, header included) formatted file containing all LEF loading sites in the following columns:
  - *name:* identifier of the loading site;
  - *position [integer≥ 1]:* start position of the loading site given in bp. The corresponding bead number will be *position/resolution_factor +1* with *resolution_factor=2000* bp the resolution of the model. Note that *position* represents the position along the simulated chain and not necessarily the actual genomic position ;
  - *length [integer≥ 1]:* length of the loading site in bp;
  - *factor [real≥ 0]:* site loading's strength. Then binding rate $k_b(i)$ at position *i* is given by $factor(i) \times kb/\left(\left[\sum_j factor(j)\right]/Nchain\right)$. Note that for monomer not present in the list of binding sites $factor(i)$=*basal_loading_factor* (see just below).
- *basal_loading_factor [real≥ 0]* defines the basal loading factor for monomers not present in the list of binding sites (see just above).
- *boundary* specifies a file path to a CSV (comma-separated values, header included) formatted file containing all boundary elements in the following columns:
  - *name:* identifier of the boundary;
  - *midpoint [integer≥ 1]:* position of the boundary given in bp;
  - *impermeability [− 1 ≤real≤ 1]*: impermeability of the boundary. The LEF motion rate $k_m(i, s)$ at position *i* in direction $s = \pm 1$ is given by *(1-|impermeability(i)|)km* if *sign(impermeability)=sign(s).*

- *boundary_direction [-1, 0 or 1*] defines a multiplicative factor applied to all boundaries. All impermeability parameters (see above) are modified to *impermeability*boundary_direction*.
- *zloop [true or false]* defines whether LEFs can traverse one another during extrusion. *zloop=true* refers to phantom collisions.
- *unidirectional [true or false]* defines whether LEFs extrude asymmetrically (*true*) or symmetrically (*false*).

For simulation parameters:

- *Niter [integer*$\geq 1$*]* defines the number of simulated trajectories. We recommend simulating at least 100 trajectories for good statistics.
- *Nmeas [integer*$\geq 1$*]* defines the number of snapshots that will be extracted from each trajectory.
- *Ninter [integer*$\geq 1$*]* is the number of MCS steps between two snapshots. The duration of one trajectory in real time will thus be $Niter \times (Nmeas - 1) \times 5msec$ (*Nmeas-1* because 1 snapshot is taken at time=0). We recommend simulating the system at least 2 hour to allow LEF and polymer dynamics to reach a steady-state.
- *init_mode [h or z]* defines how initial configurations of the polymer are randomly generated. Possible values are ‘h’ for a helical scaffold, and ‘z’ for a zigzag scaffold (Ghosh and Jost, 2018). We recommend using zigzag-like mode.

For analysis parameters (see also output files below):

- *radius_contact [real*$\geq 0$*]* corresponds to the maximal pairwise distance that defines a contact between two monomers. It is given in lattice unit with 1 lattice unit=70.7 nm.
- *chrom* defines the chromosome name to be used in output files (*hic, hic3d, chip*).
- *cmp_chrs* defines a list of synonyms designing the same chromosome (different naming conventions) for which experimental Hi-C data will be compared to predictions with chi2-min scores (see below).
- *exp_cool* specifies a file path to a Hi-C file in the cooler format from an experiment of interest, which will be used to compute chi2-min scores (see below).
- *hic3d_factor [integer*$\geq 0$*]* specifies whether to output hic3d matrix and with what resolution reduction factor, when given and bigger than zero.
- *tads_boundary* specifies a file path in the BED (Browser Extensible Data; tab separated values, header excluded) format containing all TAD regions (with the following columns: chromosome, start position, end position, name identifier) , which will be used to compute chi2-min scores (see below).

For scheduling a series of simulations, one can use the built-in feature called 'grid' simulation (see '*3dpolys_le_runner grid_nlef_km*' command) to vary the LEFs velocity (parameter *km*) and occupancy (parameter *Nlef*). In such a way, the py3DPolyS-LE will collect all comparative statistics (see Data analysis) in a single CSV file and visualize them in a heat map (see *plot_sim_stats* command), allowing to identify the local minimum. Reasonable value range for the LEF extrusion speed would be between *10kb/min* and *120 kb/min* and for LEFs occupancy between one LEF at every 400kb and one every 40 kb).

For each simulation shown in this work (Fig.1 & Supplementary Fig.1), we used parameters *Nchain=3000, L=11*, *Niter=250, Nmeas=3, Ninter=540 000* for a total trajectory length of 1.5 h. *radius_contact=3.57*$\simeq 250nm$. Default parameters were taken for *kint, kb, ku*.

#### Output files

Snapshots are saved into the file *xyzconfig_*.out* as $N_{chain} \times 3$ matrices (one column per coordinate, one raw per monomer) one after the other for every trajectory. Monomer positions are given relatively to the position of the first monomer (taken as the origin of coordinates).
From all the snapshots taken at a given time-step *t*, the contact frequency $P_{i,j}$ between any pair *(i,j)* of monomers defined as the probability to observe *i* and *j* at a Euclidean distance less than a cutoff value $r_c$(parameter $radius\_contact$) is computed using the formula $P_{i,j}(t) = \frac{\sum_{n=1}^{Niter} integer\left(((x_i^n(t)-x_j^n(t))^2+(y_i^n(t)-y_j^n(t))^2+(z_i^n(t)-z_j^n(t))^2)<=r_c^2\right)}{Niter}$, with $(x_i^n(t), y_i^n(t), z_i^n(t))$ the coordinates of monomer *i* at time step *k* from trajectory *n*. $P_{i,j}(t)$ could be seen as a predicted Hi-C map. It is then saved as 2D matrices in the HDF5 format (dataset '/hic_map') in the files *hic_*.hdf5*. Note that a HDF5 to cooler python code is also available in our package. Optionally in a similar way, a predicted Hi-C map for triple-wise contacts is computed and saved in the files *hic3d_*.cool3d* (see "Hi-C3D for triple-wise interactions"). We also estimated, for every time-step, $C_i$ the probability to find a LEF leg bound to monomer *i* that can be seen as a virtual ChIP-seq profile. It is saved in *bedGraph* format (with columns: chromosome, start, end, value) in the files chip_lef_*.bedGraph.

### Data analysis

#### Chi2-analysis

To quantitatively compare predicted intra-chromosomal organization with experiments, we developed chi-squared-based scores that allow to compare the HiC-map ($P_{i,j}$) predicted for the last time-step and the experimental ($F_{i,j}$) Hi-C data.

For a given set of non-overlapping genomic regions (TADs for example), we first computed, for each domain *d,* the average intra-domain contact frequency as a function of the genomic distance $g$, the so-called contact decay-plot, from the predicted ($P_d(g) \equiv \left[\sum_{i=b_m(d)}^{b_M(d)-g} P_{i,i+g}\right] / \left[b_M(d) - b_m(d) - g + 1\right]$) and experimental ($F_d(g) \equiv \left[\sum_{i=b_m(d)}^{b_M(d)-g} F_{i,i+g}\right] / \left[b_M(d) - b_m(d) - g + 1\right]$) Hi-C maps where $[b_m(d) : b_M(d)]$ defined the genomic coordinates of *d.* We can then estimated for *d* a chi-squared distance

$$\chi_d^{\ 2} = \sum_{x=min\_x}^{max\_x} \frac{\left(\alpha P_d\left(10^x\right) - F_d\left(10^x\right)\right)^2}{2\sigma_d^2\left(10^x\right)}$$

where we used logarithmically-evenly-spaced genomic distance $(10^x = g)$ to describe equally-well all genomic ranges. $min\_x$ and $max\_x$ define the interval of genomic distances for which we want to compare interactions, α is a global proportional factor between predicted probabilities and experimental frequencies, $\sigma_d^2(g)$ is the standard error of $F_d(g)$.

By summing the chi-squared statistics of all domains, we define a global score $\chi^{\ 2}(\alpha) = \sum_d \chi_d^{\ 2}$ that depends on the meta-parameter α. α is determined by minimizing $\chi^{\ 2}(\alpha)$. Analytically solving $\frac{\partial \chi^2}{\partial \alpha} = 0$, we finally obtained a normalized optimal chi-squared score that

$$\chi^2_{min} = \frac{1}{2n}\left\{\left(\sum_d \sum_x \frac{F_d^2\left(10^x\right)}{\sigma_d^2\left(10^x\right)}\right) - \left(\frac{\left[\Sigma_d \Sigma_x \frac{P_d\left(10^x\right) F_d\left(10^x\right)}{\sigma_d^2\left(10^x\right)}\right]^2}{\Sigma_d \Sigma_x \frac{P_d^2\left(10^x\right)}{\sigma_d^2\left(10^x\right)}}\right)\right\}$$

where $n$ is the number of terms in the double sum $\Sigma_d \Sigma_x$[...].

This analysis is done in the Python code (see below), in the step called 'stats'. $\chi^2_{min}$ values are stored in a file sim_stats.csv, which is also used to plot the chi2-heat-map (like Fig.1B).

In Fig.1C,D of the main text, we chose $10^{min\_x} = 40$ kb and $10^{max\_x} = 4000$ kb, to investigate genomic ranges where the model converges rapidly to a metastable state and is expected to be quantitative (Ghosh and Jost, 2018). Note that the largest scales may strongly depend on the initial large-scale organization that, in the current implementation of 3DPolyS-LE, cannot be modified. Therefore great care should be taken when comparing quantitatively predicted and experimental Hi-C data for genomic distances > 10 Mbp. $\chi^2_{min}$ was computed using the list of 300 kb- and 600 kb-wide TADs used in the simulations and, as the target experimental Hi-C map, a meta-TAD-like Hi-C map $F_{i,j}$ computed from the GM12878 Hi-C data measured by (Rao *et al.*, 2014). More precisely, from the GM12878 data, we first computed the average contact frequency $P_{intra,300}(s)$ and $P_{intra,600}(s)$ as a function of the genomic distance s inside TADs of size 300±30 kb and 600±60 kb, respectively, and that harbor a corner peak (list of peaks, TADs and Hi-C data taken from (Rao *et al.*, 2014)). We also estimated the average contact frequency $P_{inter}(s)$ between loci in different TADs. Then we build $F_{i,j}$ as $F_{i,j} = P_{intra,300}(|j - i|)$ if i and j are in the same 300 kb-wide TAD, $= P_{intra,600}(|j - i|)$ if i and j are in the same 300 kb-wide TAD, $=$ $P_{inter}(|j - i|)$ if i and j belong to different TADs.

#### Hi-C3D for triple-wise interactions

By using long-read sequencing technologies (Nanopore, PaCBio), it is now possible to extract experimentally triple-wise interactions. We developed an efficient method for storing Hi-C matrices of triple-wise interactions, called here Hi-C3D, while analyzing a snapshot from a simulation and extracting pairwise interactions. The file format is similar to the cooler format (Abdennur and Mirny, 2019) for a Hi-C matrix of pairwise interactions, but with an additional bins/*bin3_id* dataset for the third coordinate. Similarly to pair-wise contacts, the contact frequency $P_{i,j,k}(t)$ between any triple *(i,j,k)* of monomers at time-step *t* is defined as the probability to observe *i, j,* and *k* at a Euclidean distance less than a cutoff value $r_c$(parameter *radius_contact*) and is computed using the formula $P_{i,j,k}(t) = \frac{\sum_{i=1}^{Niter} integer\left(F^n_{i,j}(t)\ \&\ F^n_{i,k}(t)\ \&\ F^n_{k,j}(t)\right)}{Niter}$, with

$F^n_{i,j}(t) = \left[((x_i^n(t) - x_j^n(t))^2 + (y_i^n(t) - y_j^n(t))^2 + (z_i^n(t) - z_j^n(t))^2) <= r_c^2\right]$. Extracting such triple-wise interactions without optimization could be heavily computational. We thus developed an algorithm (Supplementary Figure 2) by which in a single run throughout all the combinations of triple-wise genomic coordinates for all replica polymers with at least six times reduced memory and $O(Nchain^3)$ complexity for building the result matrix. Briefly, the algorithm performs a desired resolution reduction given by a factor (*hic3d_factor* in the code), which allows to reduce memory usage and increases the output signal. The filling of the Hi-C3D data matrix is based on one main loop throughout all measurements and three inner loops over the genomic coordinates, in our case monomer index, and updating a pre-allocated memory for the result *hic3d* matrix. To calculate the *hic3d* pixel's indexes, we use the formula $hic3d_{idx}(x, y, z, N) = \sum_{i=1}^{x-1} \sum_{j=j+1}^{N-1} \sum_{k=j+1}^{N} 1 + \sum_{j=x+1}^{y-1} \sum_{k=j+1}^{N} 1 + \sum_{k=y+1}^{z} 1 = sum_{xNN}(x, N) + sum_{yN}(y, N) + sum_z(z, y)$, which by using Faulhaber's formulas ( $\sum_{k=1}^{n} k = \frac{n(n+1)}{2}$, $\sum_{k=1}^{n} k^2 = \frac{n(n+1)(2n+1)}{6}$ ) can be simplified to the formulas used in the algorithm (Supplementary Figure 2). When reconstructing all triple-wise contact, one has to consider that the saved contacts in the *hic3d* matrix are only for one combination of X,Y,Z coordinates, where normally X<Y<Z. Thus, for representing contacts in a 3D cube, one must add all coordinate combinations (XYZ, XZY, YXZ, YZX, ZXY, ZYX) with the same number of contacts. For example, a GIF animation of the different projections *hic3D(x,:,:)* for the example shown in Fig.1D of the main text is given as a Supplementary Video.

```
# convert coordinates to index, 1-based array indexing
hic3d_idx <- function(x, y, z, N){
  sum_xNN = (x-1)*(N-1)*N - N*(x-1)*x/2 - (x-1)*(N-1)*N/2 + x*(x-1)*(x+1)/6
  sum_yN = (y-x-1)*N -((y-1)*y - (x+1)*x)/2
  sum_z = (z-y)
  return(sum_xNN + sum_yN + sum_z)
}
# check if all three loci with genomic coordinates x,y, and z are in proximity r
check_proximity <- function(polymer_pos3d, x, y, z, r){
  return((euclidean_distance(polymer_pos3d[x,], polymer_pos3d[y,]) <= r)
        & (euclidean_distance(polymer_pos3d[y,], polymer_pos3d[z,]) <= r)
        & (euclidean_distance(polymer_pos3d[x,], polymer_pos3d[z,]) <= r))
}
# extract hic3d table in a cooler like format
extract_hic3d <- function(Nchain, bin_factor, polymers_pos3d, r){
```

```
  # Nchain : polymer length in monomers
  # bin_factor : binning factor to reduce resolution
  # polymers_pos3d : 3D position configuration of all replica polymer trajectories
  # r : contact radius to check for proximity
  hic3d_N = Nchain / bin_factor # reduced resolution of the polymer length
  hic3d_len = hic3d_idx(hic3d_N - 2, hic3d_N - 1, hic3d_N, hic3d_N)
  allocate(hic3d_cool(hic3d_len, 4)) # list of size hic3d_len and 4 columns
  hic3d_cool = 0  # set all to zero
  Niter = length(polymers_pos3d) # number or replica polymer trajectories
  for (i in (1:Niter)) {
    for(x in 1:(N-2)){
      for(y in (x+1):(N-1)){
       for(z in (y+1):N) {
         if (check_proximity(polymers_pos3d[i], x, y, z, r)) {
           bin1_id = (x - 1) / bin_factor + 1
           bin2_id = (y - 1) / bin_factor + 1
           bin3_id = (z - 1) / bin_factor + 1
           hic3d_i = hic3d_idx(bin1_id, bin2_id, bin3_id, hic3d_N)
           hic3d_cool[hic3d_i, 1] = bin1_id
           hic3d_cool[hic3d_i, 2] = bin2_id
           hic3d_cool[hic3d_i, 3] = bin3_id
           hic3d_cool[hic3d_i, 4] = hic3d_cool[hic3d_i, 4] + 1
        }}}}
  }
  # filter out zero contact pixels
  hic3d_cool = hic3d_cool[hic3d_cool[, 4] != 0,]
  return(hic3d_cool)
}
```

*Supplementary figure S2: hic3d Algorithm pseudo code as an R like script*

#### Documentation

(Source: https://gitlab.com/togop/3DPolyS-LE/-/blob/develop/README.md )

### 3DPolyS-LE

3D Polymer Simulation of chromosome folding by modeled loop extrusion, boundary elements and loading sites.

### Installation

#### Requirements

Packages and libraries:

- **git** client version 2.17.1, only if you use git command to download the repository;
- **gcc** compiler version 7.5.0 or higher;
- **gfortran** compiler version 7.5.0 or higher;
- **MPI** implementation like MPICH and libmpich-dev (Debian/Ubuntu) or openMPI;
- **HDF5** libraries. Debian/Ubuntu: libhdf5-103 libhdf5-cpp-103 libhdf5-dev libhdf5-mpich-dev;
- **GNU make** version 3.81 or higher;
- **CMake** version 3.15.0 or higher;
- **Python** 3.7 or higher, all required packages are listed in the requirements.txt file and alternatively in the environment.yml file;
- **Conda** version 4.8.2 or higher.

Make sure you have installed or loaded (for HPC) above required libraries.
Typically, *gfortran* is part of *gcc*. If missed, on an HPC cluster you can check if available and load the latest version:

```
module avail gcc
module load gcc/8.2.0
```

On a Ubuntu/Debian Linux it can be installed like this:

```
sudo apt-get install gfortran
# or
sudo apt-get install gcc
```

For example, on an HPC cluster (i.e. Slurm) you might need to load the following modules:

*Conda (https://conda.io)*

```
module load Anaconda3
```

Alternative could be installation of Miniconda (https://docs.conda.io/en/latest/miniconda.html).

*HDF5 (https://www.hdfgroup.org/solutions/hdf5/)*

```
module load HDF5
```

Alternatively, on an Ubuntu/Debian Linux could be installed like hits:

```
conda install hdf5
```

*MPI (Message Passing Interface)*

```
# OpenMPI (https://www.open-mpi.org/)
module load OpenMPI

# or MPICH (https://www.mpich.org/)
module load mvapich2
```

Depending on HPC modules' availability, module names could be different. Check with:

```
module avail
```

On an Ubuntu Linux, MPI could be installed like hits:

```
sudo apt-get install openmpi-bin
# or
sudo apt-get install mpich
```

*CMake ([https://cmake.org/](https://cmake.org/))*

```
module load CMake
```

Alternatively, you can install it using Conda:

```
conda install cmake
```

The CMake module is required only when you build and install the *py3DPolyS-LE* package.

Be aware that all required HPC modules have to be loaded before you run simulations.

#### 1. Clone repository

from the master branch:

`git clone https://gitlab.com/togop/3DPolyS-LE.git`

or from the development branch:

`git clone https://gitlab.com/togop/3DPolyS-LE.git -b develop`

#### 2. Build and install

To build and install as Python package, run the following commands:

```
# go to the cloned repository project folder
cd 3DPolyS-LE
make all
```

#### Troubleshooting

Depending on your installation environment, you might want to create a dedicated Python environment.

Go to the cloned repository project's folder:

```
cd 3DPolyS-LE
```

Build the default *3DPolyS-LE*'s Python environment *py3dpolys_le*:

```
make env
```

If your default Python version is a bit old you might need to specify a newer version.
In this case, you can install the *py3dpolys_le* like that:

```
conda env create -f environment.yml python=3.9
# or
conda env create -f requirements_dev.txt python=3.8
```

Activate your *py3dpolys_le* environment:

```
conda activate py3dpolys_le
# or
source activate py3dpolys_le
```

Finally, build and install:

```
make all
```

### Usage

To run a simulation:

Create a copy of an input.cfg file and update the parameters you want.
An example copy of such a configuration file you can find in the package:
https://gitlab.com/togop/3DPolyS-LE/-/blob/master/py3dpolys_le/data/ce/input.cfg
All simulation's parameters are under section _[3dpolys*le]*, here is an example:

```
[3dpolys_le]
# default 3dpolys_le parameters' values
# polymer characteristics
Nchain = 8860
L = 16
Ea = 0.
init_mode = z

# measurements
Niter = 250
Nmeas = 3
Ninter = 840000
burnin = 0
burnout = 0
burnoutM = 0

# Loop-Extrusion factors
kb = 2.8e-6
ku = 2e-6
km = 2.7e-3
Nlef = 200
# optional lef_loading_sites.csv: name,position,length,factor
lef_loading_sites                                                          =
```

```
py3dpolys_le/data/ce/dcc_rex-sites_Crane2015_bindings.csv
basal_loading_factor = 0.
#                         optional                         boundaries.csv:
name,midpoint,impermeability,score,b-position,strand
boundary = py3dpolys_le/data/ce/dcc_mex-sites_boundaries.csv
boundary_direction = 0
z_loop = true
unidirectional = false

# analysis: experiments in silico:
# 1.42 = 100nm
radius_contact = 2.84
chrom = chrX
# [integer]: if >=1, triggers hic3d with a resolution 2000*hic3d_factor
# hic3d_factor = 10

# hic-chi2-min:
cmp_chrs=chrX,X,6
exp_cool=./test/data/wt_N2_Moushumi2020_HIC1_5000.cool
tads_boundary=./py3dpolys_le/data/ce/tad_boundaries/N2.chrX.allValidPairs.
hic.5-10kbLoops.bed
```

Be aware to update properly the *[jobrunner]* section according to your system environment.

For Slurm environment, you can use such a configuration (also could be found in the example input.cfg):

```
[job_runner]
cmd_run=shell
jobid_re=\d+$
cmd_job_dependency=--dependency=afterany:{jobid}
cmd_prefix=sbatch --job-name=3dpolys_le --time=1-00:00:00 --mem-per-cpu=8G
```

```
--nodes=1 --ntasks-per-node=1 --cpus-per-task=8 {cmd_job_dependency}

[job_runner_sim]
cmd_prefix=sbatch --job-name=sim_3dpolys_le --time=3-00:00:00
--mem-per-cpu=6G --nodes=1 --ntasks-per-node=50 --cpus-per-task=1
{cmd_job_dependency}

[job_runner_analysis]
cmd_prefix=sbatch --job-name=anl_3dpolys_le --time=1-00:00:00
--mem-per-cpu=16G --nodes=1 --ntasks-per-node=1 --cpus-per-task=4
{cmd_job_dependency}

[job_runner_stats]
cmd_prefix=sbatch --job-name=sts_3dpolys_le --time=1-00:00:00
--mem-per-cpu=16G --nodes=1 --ntasks-per-node=1 --cpus-per-task=4
{cmd_job_dependency}
```

Configuration file sections:

- *3dpolys_le* comprise all parameters for running simulations and data analysis (see above).
- *job_runner* comprise general parameters for scheduling simulation pipeline steps:
    - *cmd_run* defines how to treat the generated steps commands with valid values:
        - *shell* : execute commands
        - *stdout* : print out commands to the standard output
        - *file:<file path>* : save commands into a file. It can be overwritten by a passed *3dpolys_le_runner's --cmd_run_file* argument.
    - *jobid_re* defines a regular expression to extract a batch job identifier out of an HPC batch runner output.
    - *cmd_job_dependency* defines a template to add an HPC batch job dependency, where *{jobid}* is a placeholder for the dependency job extracted by the *jobid_re* regular expression.

    - *cmd_prefix* defines the HPC batch command prefix to be used for starting a job, where {cmd_job_dependency} is a placeholder for dependency jobs as built by the *cmd_job_dependency* parameter.
- *job_runner_sim* contains general parameters for scheduling the first model simulation step:
    - *cmd_prefix* same as in the [*job_runner*] section but specific for this kind of jobs.
- *job_runner_analysis* contains general parameters for scheduling simulation output data analysis HPC batch jobs for generating predicted Chip/HiC/HiC3D data:
    - *cmd_prefix* same as in the [*job_runner*] section but specific for such kind of HPC batch jobs.
- *job_runner_stats* contains general parameters for scheduling the simulation output data analysis HPC batch jobs for calculating *hic-hic2-min* score and generating additional plots:
    - *cmd_prefix* same as in the *[job_runner]* section but specific for such kind of HPC batch jobs.
    - *plot_format* defines plot output format. Possible values supported by Python's *matplotlib* like: *png, tif, svg*.
    - *plot_cmap* defines plotting color pallet. Possible values supported by Python's *matplotlib* like: *YlGnBu, cool, hot_r, gist_heat_r, afmhot_r, YlOrRd, Greys, gist_yarg.*

To run each step separately on a personal computer without utilizing an HPC batch system, you can use such *[job_runner*]* configuration:

```
[job_runner]
cmd_run=stdout

[job_runner_sim]

[job_runner_analysis]

[job_runner_stats]
```

With such a configuration file, you can run the steps separately like this:

running a simulation:

```
3dpolys_le -o:/path/to/sim_output_folder /path/to/input.cfg
```

running analyse step: generating predicted Chip, HiC/HiC3D

```
3dpolys_le -o:/path/to/sim_output_folder -a:/path/to/analyse_output_folder /path/to/input.cfg
```

running predicted data analyse step, calculating hic-chi2-min score and storing it into a file /path/to/sim_stats.csv:

```
3dpolys_le_stats -o /path/to/sim_output_folder -a /path/to/analyse_output_folder -i /path/to/input.cfg -f /path/to/sim_stats.csv
```

generating additional plots for contact-decay, comparing simulation with an experimental HiC data:

```
3dpolys_le_runner multi_decay_plot -o /paht/to/sim_output_folder -a /path/to/analyse_output_folder -i /path/to/input.cfg
```

Alternatively, you can use a singularity image [https://cloud.sylabs.io/library/todor/default/py3dpolys_le](https://cloud.sylabs.io/library/todor/default/py3dpolys_le) to run the above commands.

```
singularity pull library://todor/default/py3dpolys_le:latest
```

Afterwards, you can run the above commands using the following prefix:

```
singularity exec -H $HOME py3dpolys_le_latest.sif <my 3dpolys_le command>
```

Additionally, you can add a batch command prefix to run it in your HPC like IBM's LSF for example:

```
bsub -n 12 -R "rusage[mem=8192]"
```

Demo data and example configuration can be found here https://gitlab.com/togop/3DPolyS-LE/-/blob/master/test/demo_run_shell.cfg and the corresponding commands to run the demo https://gitlab.com/togop/3DPolyS-LE/-/blob/master/test/demo_run_commads.txt (update paths accordingly to your environment) with the needed data files in https://gitlab.com/togop/3DPolyS-LE/-/tree/master/test/data .

Be aware that only the single steps are supported by the singularity image for now, and NOT the scenario for running a '*3dpolys_le_runner run*' command (see below).

With a properly configured *input.cfg* file for your HPC (so far tested only on Slurm) you can start a simulation job including all the above steps with the following command:

```
3dpolys_le_runner run -i my_sim_input.cfg -o ./my_sim_out
```

It will start a series of commands including simulation, analysis, and downstream statistical analysis (hic-chi2-min score) and plots (hic, contact-decay).

It is also helpful to save the output of the main 'run' command as it will print out all executed commands and in case of some errors you can rerun the failed one. One way to do that is to save the output in a file:

```
3dpolys_le_runner run -i my_sim_input.cfg -o ./my_sim_out &> 3dpolys_le_runner.log
```

In case you want to generate a shell script and execute the single steps one by one, you can generate the run shell script by:

```
3dpolys_le_runner run -i my_sim_input.cfg -o ./my_sim_out --cmd_run_file run_my_sim.sh
```

To see all supported parameters, run the following command:

```
3dpolys_le_runner --help
```

For using the Python wrapper, triggering data analysis (predicted ChIP, HiC/HiC3D, chi2-min score) steps afterwards.
Alternatively, the simulation engine directly is also available via:

```
3dpolys_le -h
```

Other available commands are:

```
3dpolys_le_stats --help
```

```
plot_hic --help
```

```
plot_sim_stats --help
```

```
hdf5_to_cooler --help
```

```
hic_converters --help
```
