## Supplementary figures and images for "3DPolyS-LE: an accessible simulation framework to model the interplay between chromatin and loop extrusion"

### Supplementary Video

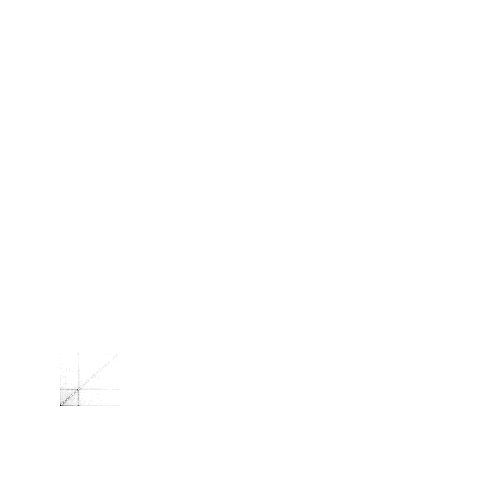
